# TaqMan Array Cards enable monitoring of diverse enteric pathogens across environmental and host reservoirs

**DOI:** 10.1101/2020.10.27.356642

**Authors:** Rachael Lappan, Rebekah Henry, Steven L. Chown, Stephen P. Luby, Ellen E. Higginson, Lamiya Bata, Thanavit Jirapanjawat, Christelle Schang, John J. Openshaw, Joanne O’Toole, Audrie Lin, Autiko Tela, Amelia Turagabeci, Tony H.F. Wong, Matthew A. French, Rebekah R. Brown, Karin Leder, Chris Greening, David McCarthy

## Abstract

**Background:** Multiple bacteria, viruses, protists, and helminths cause enteric infections that greatly impact human health and wellbeing. These enteropathogens are transmitted via several pathways through human, animal, and environmental reservoirs. Individual quantitative PCR (qPCR) assays have been extensively used to detect enteropathogens within these types of samples, whereas the TaqMan Array Card (TAC) that allows simultaneous detection of multiple enteropathogens has only previously been validated in human clinical samples.

**Methods:** Here, we performed a comprehensive double-blinded comparison of the performance of a custom TAC relative to standard qPCR for the detection of eight enteric targets, by using spiked samples, wastewater from Melbourne (Australia), and human, animal, and environmental samples from informal settlements in Suva, Fiji.

**Findings:** Both methods exhibited high and comparable specificity (TAC: 100%, qPCR: 94%), sensitivity (TAC: 92%; qPCR: 100%), and quantitation accuracy (TAC: 91%; qPCR: 99%) in non-inhibited sample matrices. PCR inhibitors substantially impacted detection via TAC, though this issue was alleviated by 10-fold sample dilution. Among samples from informal settlements, the two techniques were comparable for detection (89% agreement) and quantitation (R^2^ = 0.82). The TAC additionally included 38 other targets, enabling detection of diverse faecal pathogens and extensive environmental contamination that would be prohibitively labour intensive to assay by standard qPCR.

**Interpretation:** Overall, the two techniques produce comparable results across diverse sample types, with qPCR prioritising greater sensitivity and quantitation accuracy, and TAC trading small reductions in these for a cost-effective larger enteropathogen panel that enables a greater number of enteric pathogens to be analysed concurrently, which is beneficial given the abundance and variety of enteric pathogens in environments such as urban informal settlements. The ability to monitor multiple enteric pathogens across diverse reservoirs in turn allows better resolution of pathogen exposure pathways, and the design and monitoring of interventions to reduce pathogen load.

**Funding:** Wellcome Trust *Our Planet, Our Health* program [OPOH grant 205222/Z/16/Z].

## 1. Introduction

Diarrhoeal disease due to inadequate sanitation and poor water quality is a major public health issue and the target of one of the United Nations Sustainable Development Goals (SDG 6). This problem disproportionately affects lower- and middle-income countries, especially people living in urban informal settlements.^1,2^ Approximately 500,000 children under the age of five die from diarrhoeal disease each year,^3–5^ despite the potential to prevent an estimated 360,000 child deaths by improvements to water, sanitation and hygiene (WASH).^6^ Various nondiarrhoeal pathogens, most notably helminths, also contribute to enteric disease burden and malnutrition^7^. Moreover, asymptomatic or subclinical carriage of various enteropathogens also impacts child growth.^8^ Recent evidence has suggested that traditional household level WASH interventions such as pit latrines, handwashing with soap, and chlorination of water deliver suboptimal reductions in enteric disease in environments that are densely populated,^9^ highly contaminated^10^ or have a high prevalence of diarrhoea.^11^ This is likely due to the inability of these interventions to address the many pathways that connect environmental enteropathogens to community residents. Humans, animals, and their surrounding environments can serve as extensively interconnected reservoirs for enteropathogens. Thus, unified ‘One Health’ and ‘Planetary Health’ approaches are needed to identify pathogen exposure pathways and inform interventions that reduce pathogen load in the environment and in turn reduce human exposure.^12^

Assessing the extent of enteropathogen contamination and the impact of new mitigating interventions in urban informal settlements requires methods that can monitor several enteropathogen species in a range of sample types. Screening for a large number of enteropathogens is important, as multiple viruses, bacteria, protists and helminths are responsible for poor gastrointestinal health and diarrhoeal disease^13,14^ and mixed infections are common.^14^ Additionally, the relative contribution of individual pathogens to disease burden varies across settlements, within a settlement over time, and between individuals. Interventions can also disrupt some transmission pathways more effectively than others.^10^ The catch-all approach has traditionally been challenging due to the large number of possible enteropathogens, with a simpler solution to rely on bacterial indicator organisms to identify faecal contamination.^15,16^ However, faecal indicators do not correlate well with pathogen abundance and distribution,^17–20^ and reliance on indicators misses the complexities of enteropathogen diversity and pathogen-specific impacts of an intervention. Thus, the development of high-throughput molecular methods for enteropathogen screening of human, animal, and environmental samples would remove the need to rely solely on faecal indicators and can provide a comprehensive view of enteropathogen sources and diversity.

TaqMan quantitative PCR (qPCR) is a standard technique used across the human, animal, and environmental health fields to detect and quantify pathogens based on amplification of a pathogen-specific gene sequence.^19,21–23^ This technique can be readily used to quantify pathogenic bacteria, viruses, protists, and helminths *in situ*, whereas alternative approaches such as selective cultivation, amplicon sequencing, and metagenomic sequencing are variably challenging to implement for non-bacterial targets. Moreover, given the ability to multiplex qPCR reactions and use 96-well and 384-well plates to process many samples at a time, this technique is relatively efficient, cheap, and high-throughput with respect to sample numbers. However, the price and labour time for screening many samples for several pathogens can become high, given each additional pathogen target adds to the cost of reagents, sample volume used, and preparation time. This can become prohibitive for enteropathogen detection across human, animal, and environmental samples, where the number and taxonomic diversity of enteropathogens contributing to the burden of disease may be high and is often unknown.^24^

The TaqMan Array Card (TAC), manufactured by Applied Biosystems, is a microfluidic card designed to automate several TaqMan qPCR assays per sample. Originally designed for gene expression experiments, TAC has been effectively repurposed for detection of large panels of pathogens,^25^ with successful application to human faecal,^26^ blood,^27^ cerebrospinal fluid,^28^ and nasopharyngeal^29^ samples. Generally, the ability to efficiently detect large numbers of pathogens simultaneously is accompanied by a loss of sensitivity compared to standard qPCR,^30,31^ though it is highly cost effective compared to standard qPCR for the breadth of targets that can be detected. Several large multi-centre studies have used TAC to study the aetiology of diarrhoeal disease.^14,32^ However, TAC has rarely been applied to non-human samples, with only two studies to date using TAC to detect enteropathogens in food^33^ and environmental^18^ samples. The latter study showed that TAC can detect a wide range of pathogens in soil and water samples from informal settlements in Kisumu, Kenya.^18^ However, the sensitivity, specificity, and quantitation accuracy of TAC has not yet been extensively evaluated in environmental samples in relation to the gold standard of qPCR. Thus, it is currently unclear whether the technique presents a valid alternative to standard qPCR to monitor multiple enteropathogens across different reservoirs.

In this study, we designed and evaluated a custom enteropathogen TAC that detects 46 different pathogen marker and faecal indicator genes. We first comprehensively tested the specificity, sensitivity, and accuracy of standard qPCR and TAC on a set of mock samples consisting of spiked enteropathogen genomic DNA in different sample matrices varying in PCR inhibition levels. We additionally tested both techniques on wastewater samples from Melbourne (Australia) and human stool, animal scat, environmental water, potable water, and soil samples from informal settlements in Suva, Fiji, where the diversity and prevalence of enteropathogens was not previously known. Through this approach, we demonstrate that TAC enables the reliable monitoring of multiple enteropathogens across environmental and host reservoirs.

## 2. Materials and Methods

### 2.1. Spiked sample preparation and matrix testing

Two sets of mock samples were prepared by spiking synthetic gene blocks (IDT, Australia) representing eight enteropathogen genes **(Table 1; Table S1).** Dilution of gene blocks for addition to each test matrix was conducted based on conversion of the measured nanograms of resuspended amplicon to total gene copy number using the formula:

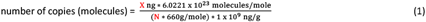

where X is the amount of measured amplicon (ng), N is the total length of dsDNA amplicon and 660 g/mole represents the average mass of 1 bp dsDNA. Set 1 consisted of 10 nuclease-free water samples for comparison of method sensitivity and specificity. Three samples contained all eight targets at low (10 copies/μl), medium (100 copies/μl), or high (1000 copies/μl) concentration; six samples contained combinations of targets and concentrations; and one sample was a blank with no targets spiked.

**Table 1.**
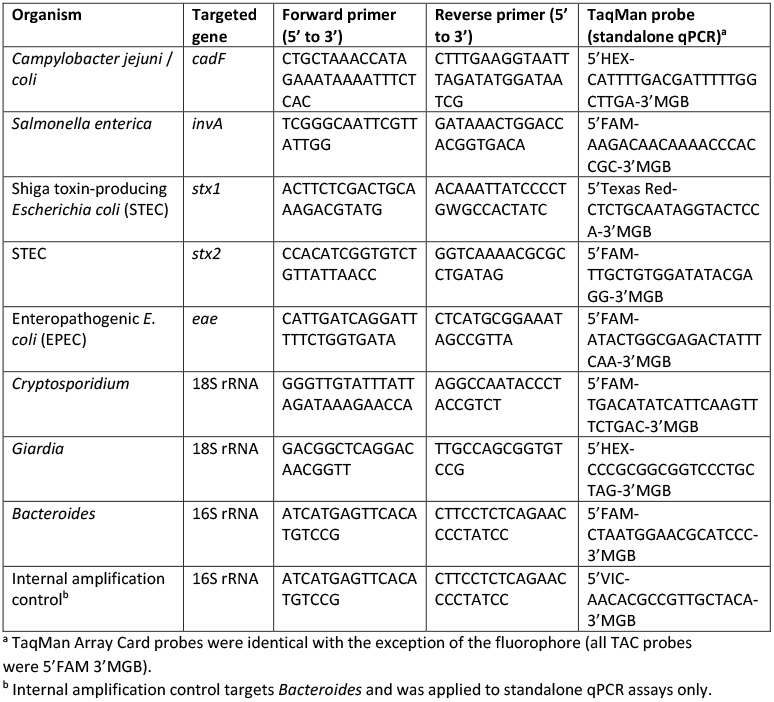
TaqMan qPCR assays used for detection by standard qPCR and custom TaqMan Array Cards.

Set 2 consisted of previously extracted Australian samples from different matrices to test the performance of the two methods under varying levels of PCR inhibition. The 36 samples tested included: nine wastewater samples with no gene blocks spiked; seven potable water samples spiked with 200 copies/μl of each target; and five different combinations of low, medium and high spiked targets in extracted DNA from each of four additional matrices (creek water, human stool, sediment and DNA extraction blank). All standard qPCR assays were performed on undiluted purified genomic DNA. As initial testing indicated that TAC reactions were inhibited for the wastewater and potable water matrices, these samples were assayed with TAC undiluted, diluted 1:10, and diluted 1:20 with nuclease-free water. Full details of the mock samples can be found in **Table S2**. All spiked samples were double-blinded and two separate laboratories delivered the results; one for TAC and another for standard qPCR.

### 2.2. Fiji sample collection and processing

Samples of child stool (n = 60), animal scats (n = 17), soil (n = 24), potable water (n = 10), and environmental water (n = 10) were collected from informal settlements in Suva, Fiji as part of the Revitalising Informal Settlements and their Environments (RISE) program (https://www.rise-program/org), a transdisciplinary research program and randomised controlled trial focused on improving environmental and human health in urban informal settlements of Fiji and Indonesia.^34,35^

Sixty child (< 5 year-old) stool samples were randomly selected for the current study from a total of 287 samples collected from 12 informal settlements during the period September 27 to November 8 2019. Samples were collected by the caregiver and stored at 4°C on frozen gel packs prior to transport to the laboratory and storage at −80°C within 24-48 hours.

Twenty water samples were collected in clean, source-water rinsed disposable bottles from the associated settlement. Potable water was run from local municipal water sources for 1 min prior to direct collection of 2 L of sample. Riverine, freshwater and stormwater (environmental water) samples were taken perpendicular from the bank and at an approximate depth of 0.15 m at each location. For each potable and environmental water sample, 1 L was filtered where possible through five 0.22 μM filters (Millipore). Where sediment prevented the passing of 1 L, a reduced volume was filtered until a total of five filters were collected. Filters were stored at −80°C within food-grade sealable bags.

Soil samples were collected using a sterile tongue depressor to transfer 2 cm^3^ of material into food-grade sealable bags. Total animal stools were collected and stored in the same manner, with visual assessment of stool age to prevent collection of older “dry” samples. Material was placed at 4°C and transferred to the laboratory within two to four hours of collection. Animal scats and soil samples were homogenised (Stomacher 400 circulator, Seward) for 1 min at 250 × rpm and stored in 0.25 g aliquots in sterile cryo-storage tubes at −80°C.

For child stool, animal scats and soil samples, total genomic DNA was isolated from 0.25 g of material using the QIAGEN DNeasy PowerSoil Pro kit as per manufacturer’s instructions, and eluted in 50 μL of sterile molecular grade water. For water samples, the filters were crushed within the bags and transferred to the bead tubes with disposable spatulas for extraction with the QIAGEN DNeasy PowerMax Soil kit with the following modifications: after the addition of buffer C1, the samples were incubated at 65°C with shaking for 30 min at 200 rpm to lyse bacterial cells. The membranes were incubated for 10 min at room temperature in 1.5 ml of nuclease-free water prior to elution in this volume. Viral RNA is co-extracted with these kits. Extracted nucleic acid samples were frozen at −80°C prior to ambient transfer and refrigeration upon receipt in Melbourne, Australia. Samples were analysed by standard qPCR and TAC within one week without refreezing.

### 2.3. Standard TaqMan qPCR detection

Standard TaqMan qPCR assays were undertaken using primers and probes for eight target pathogens **(Table 1)** under the PCR reaction and cycling conditions (40 cycles) described in US-EPA Method 1696.^36^ The PCR was conducted on a Biorad CFX96 thermocycler (Biorad, USA). Standard curves were prepared using the gene blocks **(Table S1)**^37^ serially diluted 10-fold to achieve a five-point standard curve ranging from 10^5^ to 10 copies/μL. Similarly, an internal amplification control gene block was diluted to a final concentration of 50 copies/μL with 100 copies added to each 25 μL reaction to indicate PCR inhibition.

Each 25 μL reaction contained 2 μL of either diluted standard gene block or sample genomic DNA. Reactions were conducted in triplicate for each sample and standard. Six replicates of no template controls were included on each run. The *Sketa22* assay described in Method 1696 was not performed as salmon testes DNA was not added prior to sample extraction. Quality control, data analysis and calculations were conducted as outlined in Method 1696^36^ (using https://www.epa.gov/sites/production/files/2019-04/methods-1696-1697-analysis-tool_march-2019.xltm), to ensure acceptance thresholds were met for R^2^ (standards), amplification efficiency (*E*), no-template control (NTC), method blank, internal amplification control, and lower limit of quantification (*LLOQ*). Relative fluorescence units (RFU) analysis was conducted to ensure a peak had been generated for each target assay.

### 2.4. TaqMan Array Card detection

The custom TaqMan Array Card (TAC, Applied Biosystems) contained 48 singleplex assays **(Figure S1; Table S3)**, including the eight primer and probe sets used in the standard qPCR assays and the manufacturer’s 18S rRNA control. Of the 47 custom assays, 40 have previously been validated on TAC^26,38^ and the remaining seven were assays that have previously been published as individual TaqMan qPCR assays under similar conditions.^14,39–41^ Cards were loaded with 100 μl of reaction mix per port, containing 60 μl of AgPath-ID One-Step RT-PCR master mix (Applied Biosystems; 50 μl buffer, 4 μl enzyme mix and 6 μl nuclease-free water per port) mixed with 40 μl of sample nucleic acid.^26^ Samples were diluted in nuclease-free water as necessary to allow a maximum of 1400 ng total nucleic acid per port, and 8 samples were tested per card. Loaded cards were centrifuged and sealed as per manufacturer’s instructions, and run on a QuantStudio 7 Flex instrument (Applied Biosystems) under the following cycling conditions: 45°C for 20 minutes, then 95°C for 10 minutes, followed by 45 cycles of 95°C for 15 seconds and 60°C for 1 minute.^26^

To calculate gene copies per microlitre, a standard curve was generated using synthetic plasmid controls (GeneWiz) as described in Kodani and Winchell *et al*. (2012).^42^ Three plasmids were designed with primer and probe sequences included in inserts of approximately 1 kb each (15-20 targets per plasmid). If the primers or probe were degenerate, the sequence from the reference genome was used. All three plasmid insert sequences are in **Dataset S1**. The three plasmid controls were combined at equal concentrations and seven 10-fold serial dilutions were used to make the standard curve (7.2 × 10^6^ to 7.2 copies per microlitre). This positive control was run in triplicate with a no-template control on each card. Cycle threshold (C_q_) values were manually adjusted where necessary as below, with C_q_ values exported to calculate a linear equation per target from the three replicates in R v3.6.2.^43^ Note that a standard curve could not be generated for the manufacturer’s 18S rRNA control. The lower limit of quantitation (LLOQ) was defined as the lowest dilution of the standard curve that was detectable in all three replicates. For quality control, one of the plasmids (not containing the rotavirus target) mixed with rotavirus A RNA was analysed with each new batch of master mix to test DNA polymerase and reverse transcriptase activity. A no-template control was included once every 10 cards to monitor reagent contamination.

TAC data were reviewed within the QuantStudio Real-Time PCR Software v1.3. Each multicomponent plot was manually checked for amplification, and C_q_ threshold values were checked and manually adjusted per target when the automatic threshold was inappropriate. Samples with very poor amplification curves were considered negative results, and wells flagged with both BADROX and NOISE or SPIKE, or another flag indicating a substantial issue with the well were omitted from analysis. C_q_ values were exported and analysed in R v3.6.2^43^ to calculate gene copies per microlitre of original nucleic acid extract using the standard curve for each target. For the purpose of this method comparison, all assays with a genuine amplification curve, regardless of C_q_ value, were considered positive.

### 2.5. Sensitivity and specificity analyses

Sensitivity and specificity for both methods were calculated using the spiked samples as follows:

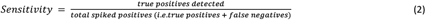

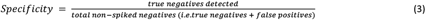

To assess quantitation accuracy, the percentage of assays that quantified gene copies per microlitre within one log_10_ of the spiked amount was calculated as follows:

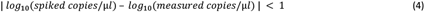

For the second set of mock samples, specificity was not calculated because it was possible for spiked targets to already be present in the samples (false positives could not be determined). Additionally, samples with a background level of target detected by either method were excluded from the quantitation accuracy calculations.

### 2.6. Statistical analysis

All statistical analyses were performed in R v3.6.2.^43^ Cohen’s κ statistic was calculated with the kappa2() function from package irr v.0.84.1^44^ to quantify agreement between standard qPCR and TAC sensitivity. Wilcoxon’s signed-rank test was applied with continuity correction using the wilcox.test() function. R^2^ values for concordance between measured qPCR and TAC gene copy numbers were calculated with the lm() function using log_10_ transformed copy numbers with a pseudocount of 1 to accommodate values of 0. Graphics were created with ggplot2 v3.3.2.^45^

## 3. Results

### 3.1. Assay sensitivity and specificity under ideal conditions

General assay sensitivity and specificity was tested using synthetic gene blocks of eight pathogen markers spiked at different concentrations into nuclease-free water in ten different combinations (80 individual assays; **Table S2 & S4**). Across all assays, the sensitivity of TAC was slightly lower (92%) than qPCR (100%). TAC performed well when detecting all eight targets at low concentration (10 copies/μl), but sometimes failed to detect targets at this concentration when others were present at high (1000 copies/μl) concentration. Specificity was very high for both assays, with no false positives detected via TAC (100%) and one false positive detected by qPCR (94%). Both methods quantified spiked targets with variable accuracy in nuclease-free water, with TAC on average underestimating target abundance by 1.73-fold and qPCR by contrast overestimated target abundance by 2.61-fold **(Figure 1; Table S4)**. Overall, 98.75% of all qPCR results within one log of the spiked concentration, compared to 91.25% from TAC **(Table 2)**. Most (5/7) of these differences from the TAC results were instances of low-copy targets that were not detected by TAC. It should be noted that the layout of TACs prevent the generation of a standard curve for each run and C_q_ thresholds are applied independently to each card. However, minimal run-to-run variation was observed amongst the plasmid controls, with controls providing very similar C_q_ values across cards; the lowest dilution, 7.2 gene copies per microlitre, was the most variable and for some targets was not consistently detected. Altogether, these findings suggest that TAC and qPCR perform comparably in inhibitor-free sample matrices.

**Figure 1.**
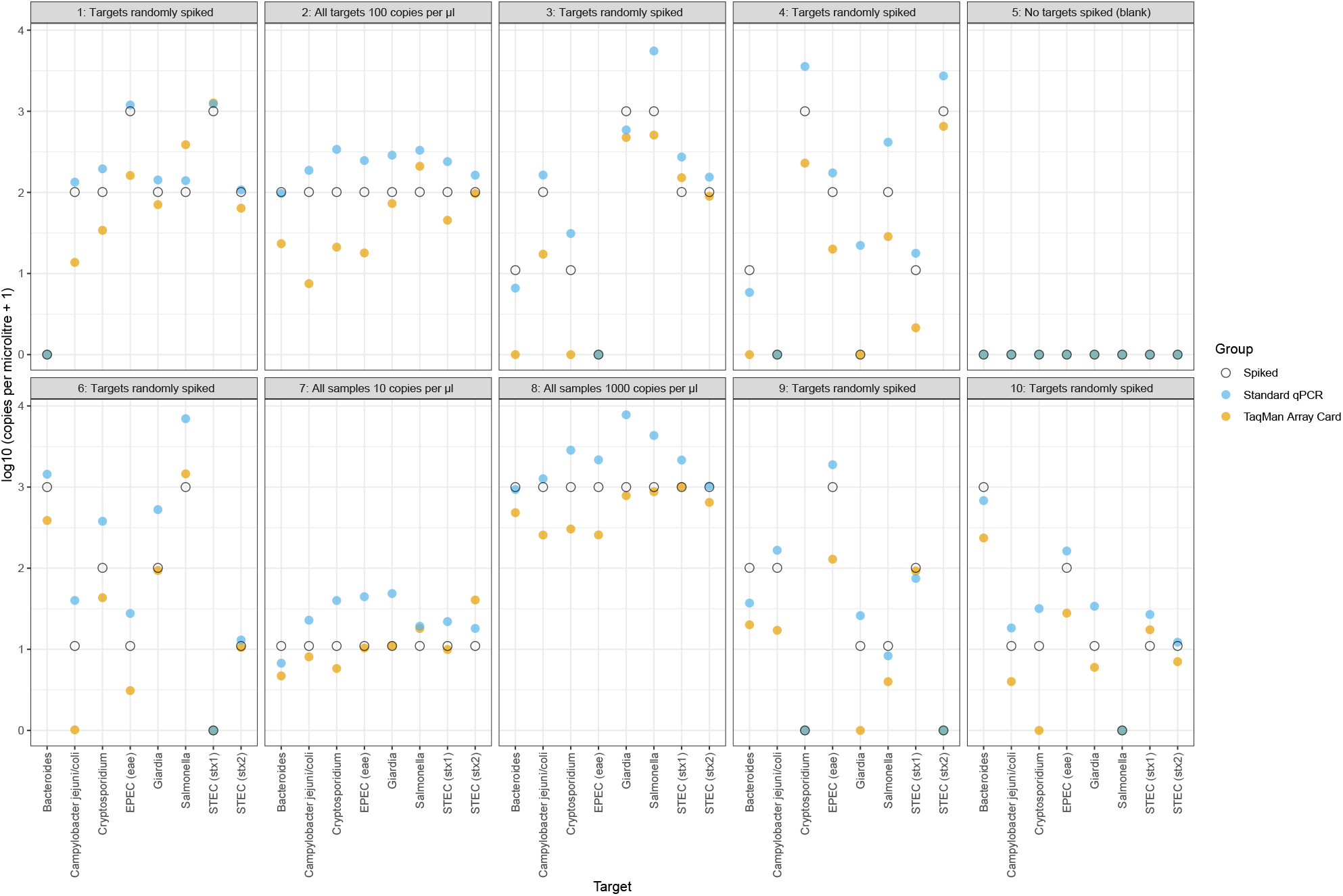
Quantitation of spiked genetic material in nuclease-free water by TAC and standard qPCR. Ten different combinations of spiked material were tested in a randomised double-blinded manner. Targets were either: spiked randomly in different combinations (samples 1, 3, 4, 6, 9, 10); spiked at consistent concentrations of 10 (sample 7), 100 (sample 2), or 1000 (sample 8) copies per microlitre; or not spiked at all (sample 5; a blank control). For each target, the quantity of material spiked (white circle), the copies detected by standard qPCR (blue circle), and the copies detected by TAC (yellow circle) are shown.

**Table 2.** Performance of TAC and qPCR on spiked samples in sample matrices varying in levels of PCR inhibitors. Results are shown by sample matrix and by target. Sensitivity is defined as the percentage of spiked targets that were detected. Accuracy is measured as percentage of assays within one log_10_ of the spiked concentration; assays where background levels of pathogen were detected by qPCR or TAC are excluded from these calculations.

| Sample matrix | Sensitivity (TAC, %) | Sensitivity (qPCR, %) | Accuracy (TAC, %) | Accuracy (qPCR, %) |
| --- | --- | --- | --- | --- |
| Nuclease-free water | 92.2 | 100 | 91.3 | 98.8 |
| Creek water | 93.3 | 96.7 | 80.0 | 80.0 |
| Sediment | 60.0 | 56.7 | 42.5 | 30.0 |
| Human stool | 83.3 | 96.7 | 80.0 | 90.0 |
| Fluorinated potable water | 71.4 | 94.6 | 62.5 | 73.2 |
| Extraction blank | 83.3 | 96.7 | 80.0 | 82.5 |
| Target |  |  |  |  |
| *Bacteroides* | 85.2 | 85.2 | 81.5 | 85.2 |
| *Campylobacter jejuni / coli* | 77.4 | 87.1 | 48.6 | 78.4 |
| *Cryptosporidium* | 77.4 | 90.3 | 70.3 | 54.1 |
| EPEC (*eae*) | 74.2 | 90.3 | 73.0 | 78.4 |
| *Giardia* | 90.3 | 96.8 | 81.5 | 81.5 |
| *Salmonella* | 83.9 | 96.8 | 78.1 | 68.8 |
| STEC (*stx1*) | 85.2 | 96.3 | 87.5 | 90.6 |
| STEC (*stx2*) | 77.4 | 93.5 | 78.4 | 91.9 |

### 3.2. Assay performance in inhibited sample matrices

The second set of test samples was used to determine the performance of each technique on samples with varying levels of PCR inhibition **(Table S2 & S4)**. For both methods, there was a reduction in sensitivity (77.3% TAC and 89.2% qPCR of spiked targets detected) and quantitation accuracy (66.7% TAC and 69.3% qPCR assays within one log of the spiked concentration) across all sample matrices **(Table 2)**. This decrease in performance compared to samples spiked in nuclease-free water **(Figure 1)** suggests both methods, especially TAC, are affected by PCR inhibitors. There was nevertheless much variability in the relative performance of the two methods across different sample matrices and pathogen targets. For example, while TAC underperformed relative to qPCR in spiked fluorinated potable water samples, the converse was true for sediment samples. Likewise, while TAC detected *Cryptosporidium* with higher accuracy, qPCR was more sensitive and accurate for detecting *Campylobacter* **(Table 2)**. TAC also detected a range of indicators and pathogens present in Melbourne sewage and stormwater samples **(Figure 3)**. TAC was more inhibited by this sample matrix than qPCR and detected no targets (including universal 16S rRNA) in five of the eight samples. However, diluting samples (1:10 and 1:20) greatly improved detection for all samples, resulting in 216-fold and 273-fold increases in the number of targets detected respectively **(Table S5)**. These results suggest that TAC is generally, though not consistently, less sensitive and accurate than qPCR for sample matrices with high inhibitor content. In common with previous findings,^18^ however, sample dilution greatly reduced inhibition without compromising detection for moderately to highly abundant targets.

### 3.3. Performance comparison with faecal and environmental samples from urban informal settlements in Fiji

A set of 121 samples from informal settlements in Fiji consisting of 60 child stool, 17 animal scats (predicted to be primarily from dogs and ducks), 20 water (10 environmental, 10 potable) samples, and 24 soil samples were analysed with TAC and standard qPCR. The nature and distribution of enteropathogen contamination in this environment is relatively unknown, and this sampling effort represents an initial insight into the baseline conditions of these settlements prior to the water and sanitation intervention to be trialled by the RISE program.^34,35^ The full dataset containing measured gene copies per microlitre is provided in **Table S6**. For the eight pathogen targets assayed by both methods, the presence/absence concordance rate was high, with 89% of all assays in agreement between both methods (Cohen’s κ = 0.619) **(Figure 2a)**. Of the remaining discordant 11%, 7% represented a detection by qPCR that was not observed with TAC, and 4% a TAC detection missed by qPCR; this indicates that the greater overall sensitivity of qPCR does not preclude the ability for TAC to detect pathogens when qPCR does not. Only one sample (a child stool) was indicated to be significantly inhibited by the qPCR *Bacteroides* internal amplification control; despite this, both methods detected the *Bacteroides* faecal indicator.

**Figure 2.**
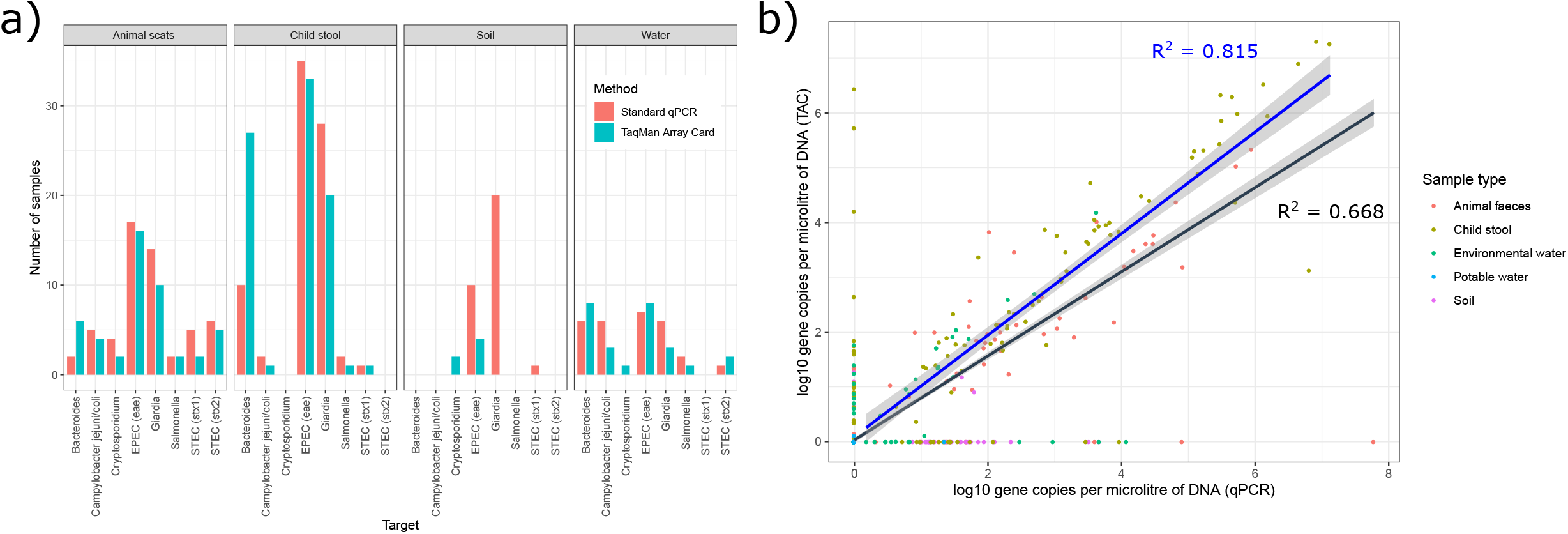
Concordance between standard qPCR and TAC in detecting pathogens in animal scats, child stool, soil, and water collected from informal settlements of Suva, Fiji. Agreement between the methods with respect to **a)** the number of positive detections of targets and **b)** the measured target quantity in log_10_ gene copies per microlitre of extracted DNA (with a pseudocount of 1 added before log_10_ transformation). The regression lines with associated 95% confidence intervals are shown for the subset of data where a target was quantified by both methods (blue, R^2^ = 0.815). Across all data points, R^2^ = 0.668 (black).

For assays where both methods detected the target, quantitation is quite consistent with R^2^ = 0.815 **(Figure 2b)**. The distribution of measured quantities for targets detected by only one method is similar on both axes, indicating that both qPCR and TAC can similarly detect targets that are missed by the other method. The target quantities measured by the two methods were significantly different (*p* = 0.00006, Wilcoxon signed rank), which was driven by instances where a target at low concentration was detected by one method and not the other **(Figure 2b)**. When considering the concordance between the techniques when a quantity was measured by both, differences in quantities were not statistically significant (*p* = 0.206, Wilcoxon signed rank).

The pathogens detected in each sample type are shown in **Figure 2a**. All targets were found in at least four samples. *Giardia* and enteropathogenic *E. coli* (EPEC; *eae* gene) were the most common of the eight targets, whereas *Salmonella* and *Cryptosporidium* were found infrequently. Reflecting the concordance rate, detections of each target in each sample type were similar. The generic faecal indicator *Bacteroides* was detected more often by TAC than by qPCR, especially in child stool samples. The major discrepancy in results was the reported detection of *Giardia* in several soil samples by qPCR (quantified at 10-100 copies per microlitre of original sample), which were not detected by TAC. It is possible that these hits are true positives that were not detected by TAC due to a combination of low *Giardia* levels, sample dilution, and challenging detection in a soil matrix. However, it is also possible that false positive detection underlies these issues given *Giardia* qPCRs accounted for the only false positive detected in the spiking study **(Figure 1)** and thus further work is required to discriminate this. Negative extraction controls were free of amplification, with the exception of a low concentration of *Giardia* in the animal scat control (detected by both methods) and in the soil control (detected only by qPCR).

### 3.4. Detection of other pathogens by TAC

In addition to the eight targets assayed via both methods, the custom TAC was designed to detect a range of other viral, bacterial, protist, and helminth enteropathogen targets **(Figure S1; Table S3)**. Of the 48 targets on the card, 44 were detected at least once (39 pathogen targets, 3 faecal indicators, 2 controls); astrovirus, *S. enterica* serovar Typhi (Typhoid fever), *Necator americanus* (hookworm), and *Cystoisospora belli* (isosporiasis) were not detected in any sample. Overall, most samples contained a wide range of enteropathogens **(Figure 3)**, with the exception of potable water which contained only very low concentrations of human faecal indicators. Environmental water and animal scat samples were the most rich in enteropathogens, with most samples containing more than eight pathogens and ten targets **(Figure 3c)**. Somewhat fewer pathogens were detected in child stool and soil (av. 2.8 and 2.2 pathogens per sample respectively, excluding faecal indicators).

**Figure 3.**
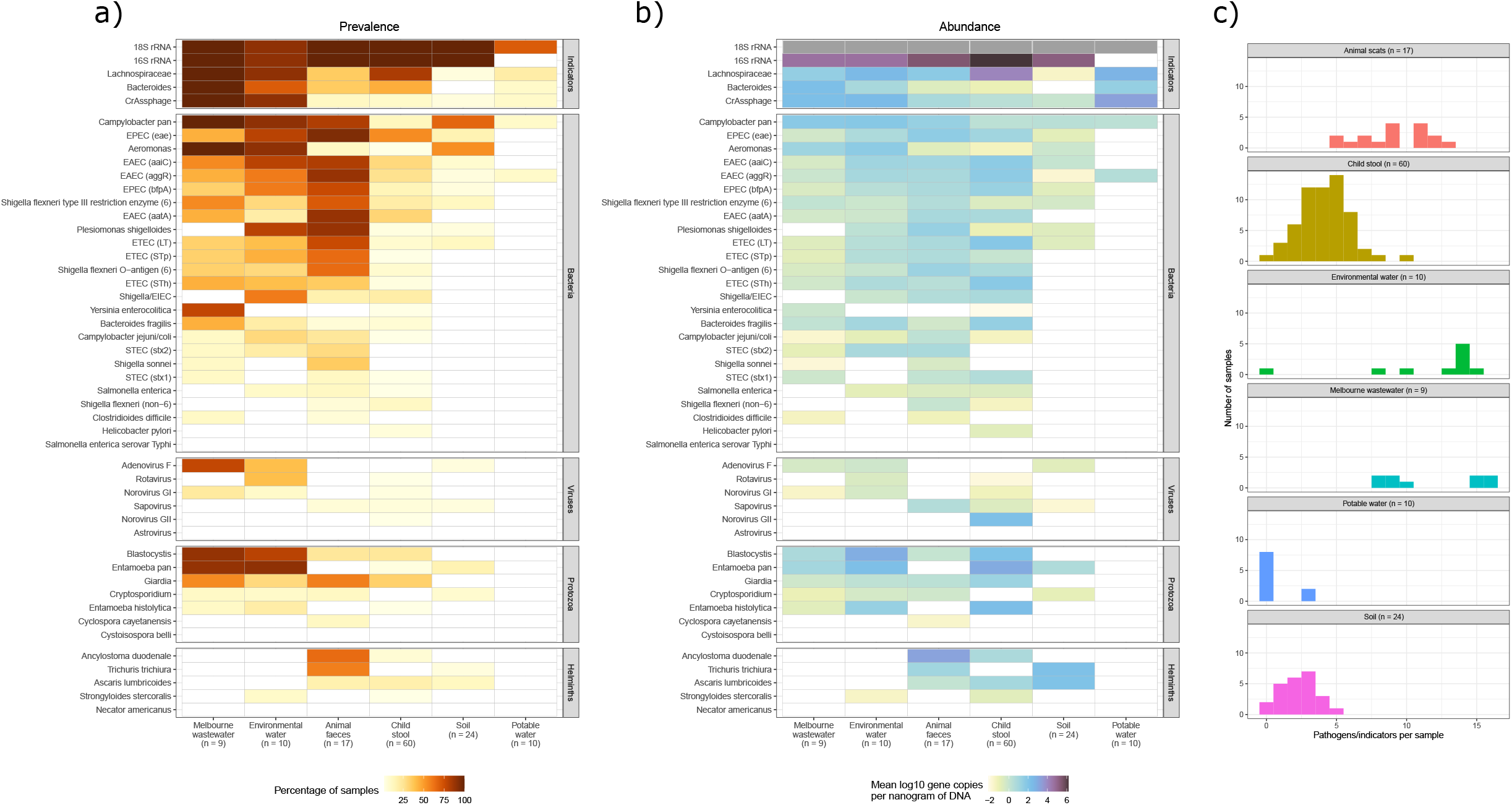
Pathogen and indicator targets detected via TaqMan Array Card in Melbourne wastewater samples and animal scats, child stool, soil, and water collected from informal settlements of Suva, Fiji. Heatmaps represent the **a)** prevalence (percentage of positive samples) and **b)** abundance (mean value of log_10_ gene copies per nanogram of DNA across positive samples) of each target by sample type. White represents a zero value, and 18S rRNA quantitation was unavailable. The number of pathogens or indicators detected per sample is represented by **c)** histograms, also by sample type. This excludes 16S rRNA and 18S rRNA and counts pathogens with multiple gene targets once.

The most prevalent enteropathogens across host and environmental reservoirs were enteroaggregative *E. coli* (EAEC) and EPEC. All three target genes were commonly detected for EAEC (*aaiC*, *aatA*, *aggR*), whereas the *eae* gene was detected more frequently than *bfpA* for EPEC (**Figure 2a)**. Both markers of *Shigella flexneri* clade 6 (O-antigen, type 3 restriction enzyme) were also present in 18 samples.^14^ *Giardia* and *Blastocystis* were common protists, with *Giardia* present at highest concentrations in human stool (**Figure 2b**). Amongst helminths, the large roundworm *Ascaris* was most common in human faecal samples, which is concordant with findings that this genus infects approximately a sixth of the world’s population.^46^ In contrast, *Ancylostoma* and *Trichuris* were predominant in animal faeces. Few helminths were detected in soil, but those that were present had moderately high abundance (approximately 120-140 copies per ng of DNA). Viruses were less prevalent overall; rotavirus and adenovirus F were most commonly detected, primarily in environmental water, whereas norovirus GII was abundant in one stool sample **(Table S6)**. Some other targets, notably *Campylobacter* pan, *Entamoeba* pan, *Aeromonas*, and *Plesiomonas shigelloides*, were abundant in environmental waters and other samples. However, they were infrequently found in child stool.

TAC also detected a range of faecal indicators and other marker genes. The universal bacterial marker 16S rRNA was detected in most samples, absent in only one environmental water and all potable water samples. Providing an estimate of total bacterial load, 16S rRNA quantities were highest in the human faecal samples and lowest in environmental water. Aside from the universal 16S rRNA assay, the most common target detected overall was human-associated *Lachnospiraceae*, a faecal marker detected in the majority of child stool and environmental water samples. This suggests that *Lachnospiraceae* is a useful faecal indicator in this population, in agreement with previous studies.^39,47^ CrAssphage, an abundant bacteriophage of human *Bacteroides* and a proposed faecal indicator,^41,48^ was the least common faecal indicator detected in individual stool samples, but was present in most of the environmental water samples. These reportedly human-specific faecal indicators were also detected in the animal scats, though to a lesser extent than in child stool **(Figure 2)**.

## 4. Discussion

Until now, the TaqMan Array Card has been validated for pathogen detection in human clinical samples,^26,30,31^ and has, to the best of our knowledge, been used in only two studies to date to detect enteropathogens in non-human samples.^18,33^ In the current study, to evaluate the performance of TAC compared to standard qPCR on environmental samples, we compared enteropathogen assays with both techniques using spiked samples of known concentration, wastewater samples from Melbourne (Australia), and a range of sample types collected from urban informal settlements in Fiji. We found that the performance of TAC in environmental samples was comparable to standard qPCR with respect to specificity, sensitivity, and quantitation accuracy in clean sample matrices. Nevertheless, TAC was somewhat less sensitive than standard qPCR in detecting spiked targets in matrices with variable inhibition. Both assays varied in quantitation accuracy depending on sample matrix and pathogen target, with TAC underestimating abundance by 1.13-fold and qPCR overestimating abundance by 1.48-fold on average across the entire dataset of spiked samples. The capacity of TAC to efficiently detect multiple enteropathogen targets potentially counterbalances these tradeoffs in sensitivity and accuracy, given the benefits of monitoring a large array of enteropathogens in samples from heavily contaminated environments. In addition, we show that TAC can effectively quantify enteric pathogens across a range of environmental, human, and animal reservoirs, thereby providing a unified method to monitor pathogen transmission pathways and evaluate public health interventions.

It can be expected that TAC would detect fewer targets per sample than qPCR. The smaller reaction volume for TAC (approximately 1 μl compared to 20 μl standard qPCRs) means there is a reduced chance that a reaction well contains a copy of a low-concentration target. Reduction in sensitivity has been observed in previous comparisons between standard qPCR and TAC. For example, Kodani *et al*. (2011) compared standard qPCR and TAC performance on respiratory specimens, and observed a general 10-fold reduction in sensitivity with TAC; some assays had a greater drop in sensitivity while others were as sensitive as standard qPCR.^49^ Subsequent studies have also reported significant reductions in sensitivity for TAC.^30,31^ Importantly, some of these previous standard qPCR/TAC comparisons have evaluated the performance of TAC relative to the standard qPCR, rather than a side-by-side comparison.^31,49^ This means that the sensitivity and accuracy of standard qPCR is assumed to be 100% (as the gold standard), whereas that for TAC is reported as the percentage of assays that agree with standard qPCR results. This may be a reasonable approach for diagnostics in clinical samples, though our tests with spiked environmental sample matrices indicated that the performance of standard qPCR is not optimal; this method overestimated target abundance overall, falsely detected *Giardia* in at least one sample, and performed suboptimally in some matrices (e.g. fluorinated water, sediment). By evaluating both methods independently, we provide a clearer view of how they each perform in challenging sample types and demonstrate that they remain comparable.

TAC was generally more susceptible to inhibition by PCR inhibitors than standard qPCR in both the spiked samples and Melbourne wastewater samples. However, sample dilution by 1:10 and 1:20 was sufficient to alleviate this and yield strong amplification curves for a wide range of indicators and pathogens. By contrast, we detected minimal inhibition in the stool, scat, soil, and water samples extracted from the Fiji informal settlements with DNeasy kits, as verified by a standard qPCR internal control. We recommend that environmental studies test for PCR inhibition prior to application of TAC to enable appropriate changes in sample preparation, such as dilution, or re-extraction of samples with optimised methods. For example, studies led by Baker ^18^ and Tsai ^33^ initially screened for inhibition with the QuantiFast pathogen kit and used 1:10 dilution to resolve it if present. Alternatively, it is possible to introduce internal amplification controls on TAC. However, we elected not to include one for this study due to the concern that the addition of control DNA would impact the detection of other targets, as enteropathogens in real environmental samples were likely to be at low concentrations and the reaction volume is minimal. Instead, we included a universal 16S rRNA target in addition to the manufacturer’s 18S rRNA control; failure of these targets to amplify indicates either substantial inhibition or minimal sample biomass, which can be discriminated by quantifying DNA through spectroscopic methods (e.g. Nanodrop, Qubit).

As both the TAC and qPCR approaches evaluated here both use the same fundamental method of TaqMan qPCR to specifically target pathogen genes of interest, both techniques are excellent options for environmental enteropathogen monitoring. As summarised in **Table 3**, they have different strengths and limitations depending on purpose of their applications. Standard qPCR offers greater sensitivity and the ease of running replicates to improve confidence in positive results and quantities, while TAC is capable of detecting up to 47 custom targets across 8 samples in a minimally laborious way, eliminating pipetting error, and greatly reducing the potential for assay contamination. It is, however, important to consider the limitations of both methods. An overarching caveat is that TAC is designed to provide the best overall result and will not be optimal for individual pathogens; optimisation of assays is more easily done for standard qPCR. Quantitation is ideally performed using a standard curve included on each run, with quality control measures associated with this.^36^ As a TAC standard curve can only occasionally be produced (requiring one whole card per replicate), the quantitation is less stringent and standard curve-related quality control measures not applicable. We demonstrated with spiked gene blocks that accurate quantitation with TAC is achievable, but the assays that target ribosomal RNA will amplify an unknown number of gene copies per organism given rRNA copy number variation.^50^ Moreover, as the protocol involves a universal reverse transcription step to detect RNA viruses, both gene and mRNA copies of all targets will be amplified. This makes it challenging to accurately estimate the number of organisms per gram or millilitre of original sample, but potentially boosts sensitivity. Some of the gene targets for bacterial pathogens assayed here are encoded on plasmids, which can be of high copy number and hence further compromise quantitation. Finally, an important consideration when interpreting qPCR-based pathogen detection is that it does not indicate organism viability. Persistence of DNA from non-viable pathogens in the environment is likely to vary by organism, and presence of DNA in stool samples without active clinical infection can also complicate interpretation of results.^51^

**Table 3.** Comparison of TAC vs qPCR for monitoring multiple pathogens.

| Method | Quantitative PCR (qPCR) and reverse transcriptase qPCR (RT-qPCR) | TaqMan Array Cards (TAC) |
| --- | --- | --- |
| Target range | **Narrow target range:** Individually detects any target of interest with appropriately optimised primers/probes. However, adding additional targets requires extra sample volume, labour, cost, and plastic waste. Assays can be multiplexed (detection of multiple targets in one reaction) with careful optimisation. | **Broad target range:** Simultaneously detects up to 47 targets and 1 internal control across 8 samples. Assays require careful card design and manufacture with appropriate lead-time. Optimisation may be required for target quantitation under universal conditions on the card. |
| Sensitivity / accuracy | **High sensitivity and accuracy:** Theoretical detection limit is approximately three gene copies per reaction. Pathogen quantitation possible with appropriate reference standards. PCR inhibition possible, but can be readily monitored with controls. | **High sensitivity / medium accuracy:** Sensitivity high but often lower than qPCR given smaller reaction volume and universal reaction conditions. Pathogen quantitation possible with reference standards, but generally requires comparisons between cards. Quantitation also challenging due to co-detection of DNA and RNA due to universal reverse transcriptase step to detect RNA viruses. Greater potential for PCR inhibition. |
| Specificity | **High specificity:** Well-designed TaqMan primer and probe sequences are very specific. | **High specificity:** Same TaqMan technology as standard qPCR. |
| Scalability | **Moderate scalability:** Extensive manual handling with large numbers of samples and/or pathogens. Large sample numbers require high labour time or robotics. Increased sample numbers require greater sample volume and produce more waste. High potential for pipetting errors. | **High scalability:** Simple and moderately fast (~3 hours) to prepare and run from extracted nucleic acids. Labour time is minimal given the few manual handling tasks required, though increases per card (eight samples). Low potential for pipetting errors. |
| Cost | **Low cost per sample:** Low reagent cost per sample (approximately USD \$2.10 for one pathogen without replicates). Small cost increase with more samples, but large increase with more targets (double the labour and reagents cost for two targets). | **Low cost per pathogen:** Moderate reagent cost per sample (approx. USD \$60). However, highly cost effective for monitoring multiple targets per sample (approx. USD \$1.28 per sample per target without replicates). |
| Resources | **Moderate resources:** Requires real-time thermal cycler. Training for molecular biology, equipment use, and software required. | **Moderate resources:** Requires real-time thermal cycler with array card block. Training needed for molecular biology, equipment, and software. |

In the context of assessing water and sanitation interventions in low-income contexts, however, a consistent method measuring relative change over time without the need to calculate organism numbers or identify aetiology is appropriate. We also included as many enteropathogen targets as possible, with no replicates. Customising cards to include fewer targets in duplicate, for example, may improve TAC sensitivity further. Liu *et al*. (2013) reported that almost half of low-concentration targets spiked into stool were detected in only one of two replicates,^26^ which is in agreement with the variability in the lowest dilution of our TAC standard curve. The feasibility of reducing the number of targets to introduce replicates is a trade-off dependent on the research project and the samples to be screened, and it may be beneficial to have some assays also available via standard qPCR to confirm ambiguous results or the presence of inhibition. Despite these limitations, we have shown TAC to perform comparably and have several advantages over standard qPCR that encompass the ability to scale up the number of targets with minimal impact on the amount of sample required, cost per sample, risk of assay contamination and error, and overall ease-of-use.

We demonstrated that there is a diverse range of enteropathogens present in urban informal settlements in Suva, Fiji, highlighting the utility of TAC in this setting. There was a high burden of bacterial pathogens in human stools, particularly EAEC and EPEC, as well as wide range of bacterial, protist, and helminth pathogens in environmental waters and animal scats. The inclusion and detection of soil-transmitted helminths in both human and environmental samples on TAC is a particularly important advance, as the standard methods for detection of such pathogens in stool involve conventional microscopy (labour-intensive, subjective, and high-risk for the operator) or serology (only viable for human-derived samples).^52^ Despite their transmission pathway, soil-transmitted helminths were more common in human and animal faeces than in soil samples from Fiji. The soil samples taken may not have been representative of helminth-contaminated areas, or alternatively detection may have been influenced by the integrity of helminth eggs impeding DNA extraction or the difficulties of detecting pathogens with low abundance in soil (demonstrated by our mock samples) combined with the small volume of soil extracted. Regarding human faecal indicators, *Lachnospiraceae* seems most suitable for this population as it was detected in the majority of faecal samples. As the assay used has been validated as highly human-specific,^39^ its presence in animal scats suggests a close association between animals and people in these settings, or possibly misidentification of scat origin. While *Bacteroides* and CrAssphage were less frequently detected in individual child stools, all three indicators were common in environmental waters. Through RISE, we are conducting further studies to understand the distribution and transmission of enteropathogens in the Fiji sites, as well as the trial settlements in Makassar, Indonesia.^35^

### Conclusion

Techniques that can adequately monitor a range of enteropathogens in humans, animals and the environment are required to assess water and sanitation improvements that aim to interrupt diverse transmission pathways. The use of qPCR in the form of individual assays or via the TaqMan Array Card enables direct detection of several enteropathogens in a range of sample types, bypassing reliance on faecal indicator organisms. Our study is the first to our knowledge to evaluate the performance of a custom TAC compared to standard qPCRs on human, animal, and environmental samples. We have demonstrated that, in these challenging sample matrices, TAC is comparable to standard qPCR and is a cost-effective, scalable, accurate, and easy to use alternative for multiple pathogens. Better understanding of the distribution, transmission, and impacts of a broad range of enteropathogens across environmental, human, and animal reservoirs is essential for improvements to public health towards SDGs 3 and 6. Among various potential applications, this will be critical for informing and evaluating future water and sanitation interventions in urban informal settlements, where the nature and extent of enteropathogen contamination is poorly characterised and diverse. More broadly, this technology enables unified approaches for surveying enteropathogens in populations and environments, as well as resolving and interrupting their transmission pathways.

## 5. Footnotes

## Acknowledgements

This work was primarily funded by the Wellcome Trust “Our Planet, Our Health” grant 205222/Z/16/Z. It was also supported by an NHMRC EL2 Fellowship (APP1178715; salary for C. G.), NHMRC Senior Research Fellowship (APP1155005; salary for K.L.) and NHMRC Equipment Grant (awarded to K.L. and S.L.C.). We also thank Eric Houpt, Darwin Operario, James Platts-Mills, Bas Dutilh, Ondrej Cinek, Brett Davis, Fiona Barker, and Sean Bay for helpful discussions. We thank David Rayner for TaqMan Array Card technical support and advice, and Sarah Thomas and Julie Bines for providing rotavirus RNA for quality control purposes. We acknowledge the hard work of the RISE field and laboratory teams in Fiji and the broader team within the RISE program. Finally, we thank all study participants in RISE.

## Author contributions

Different authors were responsible for study conception (S.L.C., K.L., S.P.L., D. M., R.R.B., T.W., M.F.), experimental design (D.M., C.G., R.H., R.L., K.L., S.L.C., S.P.L., E.H., M.F.), field sampling (R.H., D.M., J.O., J.J.O., S.P.L., A.T., A.T., C.S., A.L.), DNA extractions (C.G., T.J., L.B., R.H., D.M., C.S., E.H., A.L., J.J.O, J.O.), TaqMan array card setup and analysis (R.L., C.G., E.H., K.L., S.L.C., S.P.L., J.J.O), qPCR setup and analysis (R.H., D.M., C.S.), statistical analysis (R.L., C.G.), paper writing (R.L., C.G., R.H., D.M.), and critical review and approval of the manuscript (all authors).

## Ethics information

Ethics review and approval was provided by participating university and local IRBs, including Monash University Human Research Ethics Committee (Melbourne, Australia; protocol 9396) and the College Human Health Research Ethics Committee (CHREC) at the Fiji Institute of Pacific Health Research (FIPHR) and College of Medicine, Nursing, and Health Sciences at Fiji National University (FNU) (Suva, Fiji; protocol 137.19). All study settlements, households, and caregivers/respondents provide informed consent.

## Conflict of interest statement

The authors declare no conflict of interest.

## Notes

### Competing Interest Statement

The authors have declared no competing interest.

