## Supplementary information for "TaqMan Array Cards enable monitoring of diverse enteric pathogens across environmental and host reservoirs"

**Figure S1.** Layout of the custom TaqMan Array Card (TAC) designed and optimised for this study. Each card contains eight ports. Each port is connected to 48 wells each containing a different set of primer pairs and probes specific to the listed pathogen or indicator.

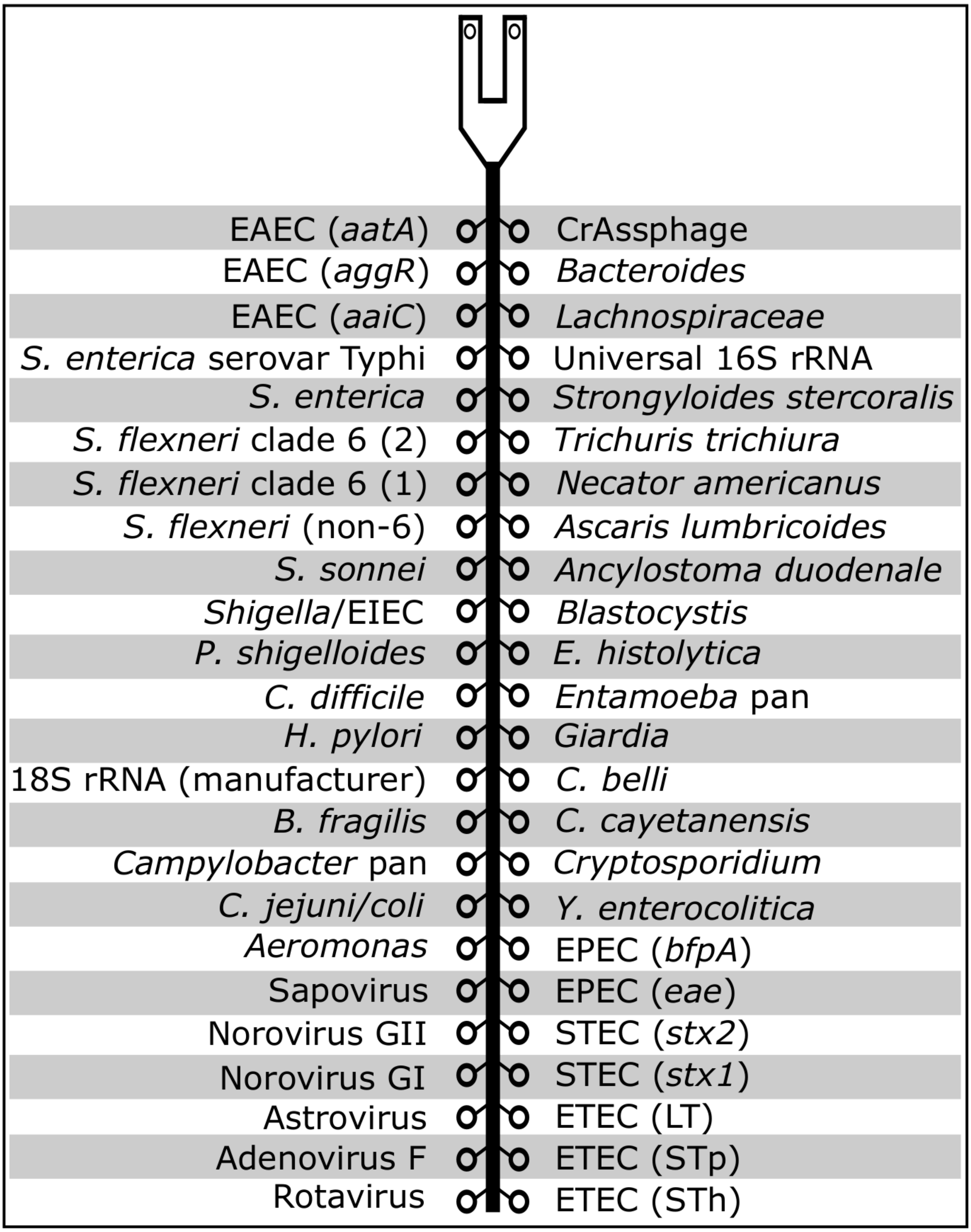

**Table S1.** Gene block fragments applied to the spiked test samples in this study.

| Organism | Target | Length (bp) | Sequence |
|---|---|---|---|
| *Campylobacter jejuni* | *cadF* | 221 | CTTTGAAGGTAATTTAGATATGGATAATCGTTATGCACCAGGGATTAGACTTGGTTATCATTTTGACGATTTTTGGCTTGATCA ATTAGAATTTGGGTTAGAGCATTATTCTGATGTTAAATATACAAATACTAATAAAACTACAGATATTACAAGAACTTATTTGAG TGCTATTAAAGGTATTGATGTAGGTGAGAAATTTTATTTCTATGGTTTAGCAG |
| *Salmonella enterica* | *invA* | 140 | GCCGATGCCGGTGAAATTATCGCCACGTTCGGGCAATTCGTTATTGGCGATAGCCTGGCGGTGGGTTTTGTTGTCTTCTCTATT GTCACCGTGGTCCAGTTTATCGTTATTACCAAAGGTTCAGAACGTGTCGCGGAAGT |
| *Escherichia coli* | *eae* | 180 | CCCGCTTTACGGCAAATTTAGGTGCGGGTCAGCGTTTTTTCCTTCCTGAAAATATGTTGGGCTATAACGTCTTCATTGATCAGGA TTTTTCTGGTGATAATACCCGTTTAGGTATTGGTGGCGAATACTGGCGAGACTATTTCAAAAGTAGTGTTAACGGCTATTTCCGC ATGAGCGGCT |
| *Escherichia coli* | *stx-1* | 132 | ACTTCTCGACTGCAAAGACGTATGTAGATTCGCTGAATGTCATTCGCTCTGCAATAGGTACTCCATTACAGACTATTTCATCAGGA GGTACGTCTTTACTGATGATTGATAGTGGCACAGGGGATAATTTGT |
| *Escherichia coli* | *stx-2* | 180 | CTCTTCGTTAAATAGTATACGGACAGAGATATCGACCCCTCTTGAACATATATCTCAGGGGACCACATCGGTGTCTGTTATTAACC ACACCCCACCGGGCAGTTATTTTGCTGTGGATATACGAGGGCTTGATGTCTATCAGGCGCGTTTTGACCATCTTCGTCTGATTATT GAGCAAAA |
| *Cryptosporidium parvum* | *18S rRNA* | 125 | GGGTTGTATTTATTAGATAAAGAACCAATTTATTGGTGACTCATAATAACTTTACGGATCACATTAAATGTGACATATCATTCAAGT TTCTGACCTATCAGCTTTAGACGGTAGGGTATTGGCCT |
| *Giardia lamblia* | *18S rRNA* | 89 | TCACCCGGGACGCGGCGGACGGCTCAGGACAACGGTTGCACCCCCCGCGGCGGTCCCTGCTAGCCGGACACCGCTGGCAACCCG GCGCC |
| *Bacteroides* spp.* | *16S rRNA* | 127 | ATCATGAGTTCACATGTCCGCATGATTAAAGGTATTTTCCGGTAGACGATGTGTAGCAACGGCGTGTTATAGTAGGCGGGGTAAC GGCCCACCTAGTCAACGATGGATAGGGGTTCTGAGAGGAAGG |
| *Bacteroides* spp | *16S rRNA* | 133 | ATCATGAGTTCACATGTCCGCATGATTAAAGGTATTTTCCGGTAGACGATGGGGATGCGTTCCATTAGCTCGAGATAGTAGGCGG GGTAACGGCCCACCTAGTCAACGATGGATAGGGGTTCTGAGAGGAAGG |

**Bacteroides* spp. internal amplification control

**Table S2 (xlsx).** Details of the samples spiked for the comparison of performance of TAC and standard qPCR. The first tab provides details of the targets spiked in nuclease-free water. The second tab provides details of the targets spiked in different matrices containing potential PCR inhibitors.

**Table S3.** List of primers and probes present in the TaqMan array card. All selected primers and probes were based on the cited references.

| **Organism** | **Target gene** | **Forward primer** | **Reverse primer** | **Probe (all 5'FAM 3'MGB)** |
|---|---|---|---|---|
| Rotavirus[1,2] | NSP3 | ACCATCTWCACRTRACCCTCTATGAG | GGTCACATAACGCCCCTATAGC | AGTTAAAAGCTAACACTGTCAAA |
| Adenovirus 40/41 (F)[2] | Fiber | AACTTTCTCTCTTAATAGACGCC | AGGGGGCTAGAAAACAAAA | CTGACACGGGCACTCT |
| Astrovirus[1,2] | Capsid | CAGTTGCTTGCTGCGTTCA | CTTGCTAGCCATCACACTTCT | CACAGAAGAGCAACTCCATCGC |
| Norovirus GI[2] | ORF1-2 | CGYTGGATGCGNTTYCATGA | CTTAGACGCCATCATCATTYAC | TGGACAGGAGATCGC |
| Norovirus GII[1,2] | ORF1-2 | CARGARBCNATGTTYAGRTGGATGAG | TCGACGCCATCTTCATTCACA | TGGGAGGGCGATCGCAATCT |
| Sapovirus[1] | RdRp | GAYCAGGCTCTCGCYACCTAC<br>TTTGAACAAGCTGTGGCATGCTAC | CCCTCCATYTCAAACACTA | CYTGGTTCATAGGTGGTRCAG<br>CAGCTGGTACATTGGTGGCAC |
| *Aeromonas*[1] | Aerolysin | TYCGYTACCAGTGGGACAAG | CCRGCAAACTGGCTCTCG | CAGTTCCAGTCCCACCACTT |
| *Campylobacter jejuni / coli*[1,2] | *cadF* | CTGCTAAACCATAGAAATAAAATTTCTCAC | CTTTGAAGGTAATTTAGATATGGATAATCG | CATTTTGACGATTTTTGGCTTGA |
| *Campylobacter* pan[2] | *cpn60* | AAAGTIGGMAAAGATGGTGTTAT<br>AAAGTIGGWAAAGACGGYGTTAT | TCAAATTGCATACCYTCAAC | TTTGCCTCTTCMACAGT<br>TTTGCTTCTTCWACAGT |
| *Bacteroides fragilis*[2] | *bft* | GGGACAAGGATTCTACCAGCTTTATA | ATTCGGCAATCTCATTCATCATT | CAATGGCGAATCCATCAG |
| *Helicobacter pylori*[2] | *ureC* | GACACCAGAAAAAGCGGCTA | AGCGCATGTCTTCGGTTAAA | TCACTAAAGCGTTTTCTACC |
| *Clostridioides difficile*[1,2] | *tcdB* | GGTATTACCTAATGCTCCAAATAG | TTTGTGCCATCATTTTCTAAGC | CCTGGTGTCCATCCTGTTTC |
| *Plesiomonas shigelloides*[2] | *gyrB* | CCGCCGTGAAGGCAAAG | GCTACCGGCTCACCCAGAT | CACACCCAAGAATAC |
| *Shigella*/EIEC[1,2] | *ipaH* | CCTTTTCCGCGTTCCTTGA | CGGAATCCGGAGGTATTGC | CGCCTTTCCGATACCGTCTCTGCA |
| *Shigella sonnei*[3] | Putative methylase | TGCCGCTAAAATCCTTCTGT | GCGTACGACGAAAGGAAAAA | GAAGTTATTGATTCCGCCC |
| *Shigella flexneri* (most serotypes except 6)[3] | Putative periplasmic protein | TGGGTGCATCCTGACCTGT | GACAAACAATAACGAGCTACCGAT | ACCACGGAATAATCCCGCAG |
| **Shigella flexneri* 6[3] | O-antigen | CTCCTATCCGTGATTATAGTGCA | GCACACACAACTCACTGTATTT | TCCTTCTCACGATTAAAATC |
| **Shigella flexneri* 6[3] | Type 3 restriction enzyme | CTTTCAACGCACGAATATCAAC | GAACCTGATCCAGACGGAGA | TTCTTCAGAACCGGGTTTTG |
| *Salmonella enterica*[1] | *invA* | TCGGGCAATTCGTTATTGG | GATAAACTGGACCACGGTGACA | AAGACAACAAAACCCACCGC |
| *Salmonella enterica* serovar Typhi[2] | *staG* (STY0201) | CGCGAAGTCAGAGTCGACATAG | AAGACCTCAACGCCGATCAC | CAGCCTGCTCCAGAACA |
| EAEC[1,2] | *aaiC* | ATTGTCCTCAGGCATTTCAC | ACGACACCCCTGATAAACAA | TAGTGCATACTCATCATTTAAG |
| EAEC[2] | *aggR* | GCAATCAGATTAARCAGCGATACA | TTCGGACAACTRCAAGCATC | AAGACGCCTAAAGGATGCCC |

| | | | | |
|---|---|---|---|---|
| EAEC[1,2] | *aatA* | CTGGCGAAAGACTGTATCAT | TTTTGCTTCATAAGCCGATAGA | TGGTTCTCATCTATTACAGACAGC |
| ETEC[1,2] | STh | GCTAAACCAGYAGRGTCTTCAAAA | CCCGGTACARGCAGGATTACAACA | TGGTCCTGAAAGCATGAA |
| ETEC[1,2] | STp | TGAATCACTTGACTCTTCAAAA | GGCAGGATTACAACAAAGTT | TGAACAACACATTTTACTGCT |
| ETEC[1,2] | LT | TTCCCACCGGATCACCAA | CAACCTTGTGGTGCATGATGA | CTTGGAGAGAAGAACCCT |
| STEC[1,2] | *stx1* | ACTTCTCGACTGCAAAGACGTATG | ACAAATTATCCCCTGWGCCACTATC | CTCTGCAATAGGTACTCCA |
| STEC[1,2] | *stx2* | CCACATCGGTGTCTGTTATTAACC | GGTCAAAACGCGCCTGATAG | TTGCTGTGGATATACGAGG |
| EPEC[1,2] | *eae* | CATTGATCAGGATTTTTCTGGTGATA | CTCATGCGGAAATAGCCGTTA | ATACTGGCGAGACTATTTCAA |
| EPEC[1,2] | *bfpA* | TGGTGCTTGCGCTTGCT | CGTTGCGCTCATTACTTCTG | CAGTCTGCGTCTGATTCCAA |
| *Yersinia enterocolitica*[2] | *lysP* | TGATTCACCAGCAGCAATAC | GGCATCATGAAAGGCGG | TGTCGGTTTCTCCTTCCAGG |
| *Cryptosporidium*[1,2] | 18S rRNA | GGGTTGTATTTATTAGATAAAGAACCA | AGGCCAATACCCTACCGTCT | TGACATATCATTCAAGTTTCTGAC |
| *Cyclospora cayetanensis*[2] | 18S rRNA | AAAAGCTCGTAGTTGGATTTCTG | AACACCAACGCACGCAGC | AAGGCCGGATGACCACGA |
| *Cystoisospora belli*[2] | ITS2 | ATATTCCCTGCAGCATGTCTGTTT | CCACACGCGTATTCCAGAGA | CAAGTTCTGCTCACGCGCTTCTGG |
| *Giardia*[1,2] | 18S rRNA | GACGGCTCAGGACAACGGTT | TTGCCAGCGGTGTCCG | CCCGCGGCGGTCCCTGCTAG |
| *Entamoeba* pan[2] | 18S rRNA | AAACGATGTCAACCAAGGATTG | TCCCCCTGAAGTCCATAAACTC | CCTTGTTCAGAACTTAAAGAGAAA |
| *Entamoeba histolytica*[1,2] | 18S rRNA | ATTGTCGTGGCATCCTAACTCA | GCGGACGGCTCATTATAACA | TCATTGAATGAATTGGCCATTT |
| *Blastocystis*[2] | 18S rRNA | TGGTCCGRTGAACACTTTGGAT | CCTACGGAAACCTTGTTACGACTTCA | CTTCCTCTAAATGRTAAGATT |
| *Ancylostoma duodenale*[2] | ITS2 | GAATGACAGCAAACTCGTTGTTG | ATACTAGCCACTGCCGAAACGT | ATCGTTTACCGACTTTAG |
| *Ascaris lumbricoides*[2] | ITS1 | GCCACATAGTAAATTGCACACAAAT | GCCTTTCTAACAAGCCCAACAT | TTGGCGGACAATTGCATGCGAT |
| *Necator americanus*[2] | ITS2 | CTGTTTGTCGAACGGTACTTGC | ATAACAGCGTGCACATGTTGC | CTGTACTACGCATTGTATAC |
| *Trichuris trichiura*[1,2] | 18S rRNA | TTGAAACGACTTGCTCATCAACTT | CTGATTCTCCGTTAACCGTTGTC | CGATGGTACGCTACGTGCTTACCATGG |
| *Strongyloides stercoralis*[2] | Dispersed repetitive sequence | TCCAGAAAAGTCTTCACTCTCCAG | TGCGTTAGAATTTAGATATTATTGTTGCT | TCAGCTCCAGTTGAACAACAGCCTCCAA |
| Universal bacterial[1] | 16S rRNA | TCCTACGGGAGGCAGCA | GGACTACCAGGGTATCTAATCCTG | CGTATTACCGCGGCTGCT |
| Lachnospiraceae (Lachno3 faecal indicator)[4] | 16S rRNA | CAACGCGAAGAACCTTACCAAA | CCCAGAGTGCCCACCTTAAAT | CTCTGACCGGTCTTTAATCGGA |
| *Bacteroides* (HF183/BacR287 faecal indicator)[5] | 16S rRNA | ATCATGAGTTCACATGTCCG | CTTCCTCTCAGAACCCCTATCC | CTAATGGAACGCATCCC |
| CrAssphage (CPQ_056 faecal indicator)[6] | orf00024 | CAGAAGTACAAACTCCTAAAAAACGTAGAG | GATGACCAATAAACAAGCCATTAGC | AATAACGATTTACGTGATGTAAC |

* Both assays required to be positive for *S. flexneri* serotype 6 to be detected, as per Liu *et al.* (2016)[3].

**Table S4 (xlsx).** Comparison of target copies quantified by TAC and standard qPCR for the spiked samples in different matrices.

**Table S5 (xlsx).** Comparison of the effects of sample dilution on target detection by TAC in sewage samples.

**Table S6 (xlsx).** Comparison of target copies quantified by TAC and standard qPCR for the human stool, animal scat, soil, and water samples from urban informal settlements of Suva, Fiji.

**Dataset S1 (docx).** Information on the three plasmid inserts used for production of standard curves. Each plasmid contains synthetic primers and probes for each pathogen / indicator target.
