## Supplementary material for "TaqMan Array Cards enable monitoring of diverse enteric pathogens across environmental and host reservoirs": Dataset S1

**Dataset S1. Information on the three plasmid inserts used for production of standard curves.** Each plasmid contains synthetic primers and probes for each pathogen / indicator target.

Primers are underlined, with probes in bold.

### **Plasmid 1 insert:**

ACCATCTACACATGACCCTCTATGAGT**AGTTAAAAGCTAACACTGTCAAA**GCTATAGGGGCGTTATGTGACCAACTTTCTCTCTTAATAGACGCCA**CTGACACGGGCACTCT**TTTTGTTTTCTAGCCCCCTCAGTTGCTTGCTGCGTTCAT**CACAGAAGAGCAACTCCATCGC**AGAAGTGTGATGGCTAGCAAGCGCTGGATGCGCTTCCATGACAAGAGCCAATGTTCAGATGGATGAG**TGGACAGGAGATCGCTGGGAGGGCGATCGCAATCT**GTAAATGATGATGGCGTCTAAGTGTGAATGAAGATGGCGTCGAGATCAGGCTCTCGCCACCTACTTTGAACAAGCTGTGGCATGCTAC**CCTGGTTCATAGGTGGTACAGCAGCTGGTACATTGGTGGCAC**TAGTGTTTGAGATGGAGGGCCGGCAAACTGGCTCTCGA**CAGTTCCAGTCCCACCACTT**CTTGTCCCACTGGTAACGAACTTTGAAGGTAATTTAGATATGGATAATCGA**CATTTTGACGATTTTTGGCTTGA**GTGAGAAATTTTATTTCTATGGTTTAGCAGTCAAATTGCATACCTTCAACG**TTTGCCTCTTCAACAGT**ATAACACCATCTTTGCCCACTTTGGGACAAGGATTCTACCAGCTTTATAG**CAATGGCGAATCCATCAG**AATGATGAATGAGATTGCCGAATGACACCAGAAAAAGCGGCTAC**TCACTAAAGCGTTTTCTACC**TTTAACCGAAGACATGCGCTTTTGTGCCATCATTTTCTAAGCA**CCTGGTGTCCATCCTGTTTC**CTATTTGGAGCATTAGGTAATACCCCGCCGTGAAGGCAAAGT**CACACCCAAGAATAC**ATCTGGGTGAGCCGGTAGCCCTTTTCCGCGTTCCTTGAC**CGCCTTTCCGATACCGTCTCTGCA**GCAATACCTCCGGATTCCGTGCCGCTAAAATCCTTCTGTA**GAAGTTATTGATTCCGCCC**TTTTTCCTTTCGTCGTACGCTGGGTGCATCCTGACCTGTC**ACCACGGAATAATCCCGCAG**ATCGGTAGCTCGTTATTGTTTGTC

**Expected hits for all primers and probes in plasmid 1 (5’ to 3’):**

Rotavirus F: 1-26
Rotavirus P: 28-50
Rotavirus R: 72-51

Adenovirus F: 73-95
Adenovirus P: 97-112
Adenovirus R: 131-113

Astrovirus F: 132-150
Astrovirus P: 152-173
Astrovirus R: 194-174

Norovirus GI F: 195-214
Norovirus GII F: 215-240
Norovirus GI P: 241-255
Norovirus GII P: 256-275
Norovirus GI R: 297-276
Norovirus GII R: 318-298

Sapovirus F(1): 319-339
Sapovirus F(2): 340-363
Sapovirus P(1): 364-384
Sapovirus P(2): 385-405
Sapovirus R: 424-406

Aeromonas F: 483-464
Aeromonas P: 444-463
Aeromonas R: 425-442

C. jejuni/coli F: 567-538
C. jejuni/coli P: 515-537
C. jejuni/coli R: 484-513

Campy F: 628-606
Campy P: 589-605
Campy R: 568-587

ETBF F: 629-654
ETBF P: 656-673
ETBF R: 696-674

H. pylori F: 697-716
H. pylori P: 718-737
H. pylori R: 757-738

C. difficile F: 824-801
C. difficile P: 781-800
C. difficile R: 758-779

P. shigelloides F: 825-841
P. shigelloides P: 843-857
P. shigelloides R: 876-858

Shig/EIEC F: 877-895
Shig/EIEC P: 897-920
Shig/EIEC R: 939-921

S. sonnei F: 940-959
S. sonnei P: 961-979
S. sonnei R: 999-980

S. flexneri periplasmic F: 1000-1018
S. flexneri periplasmic P: 1020-1039
S. flexneri periplasmic R: 1063-1040

### **Plasmid 2 insert:**

CTCCTATCCGTGATTATAGTGCAG**TCCTTCTCACGATTAAAATC**AAATACAGTGAGTTGTGTGTGCCTTTCAACGCACGAATATCAACA**TTCTTCAGAACCGGGTTTTG**TCTCCGTCTGGATCAGGTTCTCGGGCAATTCGTTATTGGC**AAGACAACAAAACCCACCGC**TGTCACCGTGGTCCAGTTTATCAAGACCTCAACGCCGATCACT**CAGCCTGCTCCAGAACA**CTATGTCGACTCTGACTTCGCGATTGTCCTCAGGCATTTCACT**TAGTGCATACTCATCATTTAAG**TTGTTTATCAGGGGTGTCGTGCAATCAGATTAAGCAGCGATACAT**AAGACGCCTAAAGGATGCCC**GATGCTTGCAGTTGTCCGAACTGGCGAAAGACTGTATCATA**TGGTTCTCATCTATTACAGACAGC**TCTATCGGCTTATGAAGCAAAAGCTAAACCAGTAGAGTCTTCAAAATGAATCACTTGACTCTTCAAAA**TGGTCCTGAAAGCATGAATGAACAACACATTTTACTGCT**AACTTTGTTGTAATCCTGCCTGTTGTAATCCTGCTTGTACCGGGTTCCCACCGGATCACCAAG**CTTGGAGAGAAGAACCCT**TCATCATGCACCACAAGGTTGACTTCTCGACTGCAAAGACGTATGG**CTCTGCAATAGGTACTCCA**GATAGTGGCACAGGGGATAATTTGTCCACATCGGTGTCTGTTATTAACCT**TTGCTGTGGATATACGAGG**CTATCAGGCGCGTTTTGACCCATTGATCAGGATTTTTCTGGTGATAA**ATACTGGCGAGACTATTTCAA**TAACGGCTATTTCCGCATGAGTGGTGCTTGCGCTTGCTC**CAGTCTGCGTCTGATTCCAA**CAGAAGTAATGAGCGCAACGTGATTCACCAGCAGCAATACA**TGTCGGTTTCTCCTTCCAGG**CCGCCTTTCATGATGCCGGGTTGTATTTATTAGATAAAGAACCAG**TGACATATCATTCAAGTTTCTGAC**AGACGGTAGGGTATTGGCCT

**Expected hits for all primers and probes on plasmid 2 (5’ to 3’)**

S. flexneri 6-O F: 1-23
S. flexneri 6-O P: 25-44
S. flexneri 6-O R: 66-45

S. flexneri 6-T3 F: 67-88
S. flexneri 6-T3 P: 90-109
S. flexneri 6-T3 R: 129-110

Salmonella F: 130-148
Salmonella P: 150-169
Salmonella R: 191-170

Typhi F: 251-230
Typhi P: 213-229
Typhi R: 192-211

aaiC F: 252-271
aaiC P: 273-294
aaiC R: 314-295

aggR F: 315-338
aggR P: 340-359
aggR R: 379-360

aatA F: 380-399
aatA P: 401-424
aatA R: 446-425

STh F: 447-470
STp F: 471-492
STh P: 493-510
STp P: 511-531
STh R: 575-552
STp R: 551-532

LT F: 576-593
LT P: 595-612
LT R: 633-613

Stx1 F: 634-657
Stx1 P: 659-677
Stx1 R: 702-678

Stx2 F: 703-726
Stx2 P: 728-746
Stx2 R: 766-747

eae F: 767-792
eae P: 794-814
eae R: 835-815

bfpA F: 836-852
bfpA P: 854-873
bfpA R: 893-874

Y. entero F: 894-913
Y. entero P: 915-934
Y. entero R: 951-935

Crypto F: 952-978
Crypto P: 980-1003
Crypto R: 1023-1004

**Plasmid 3 insert:**

AACACCAACGCACGCAGCT**AAGGCCGGATGACCACGA**CAGAAATCCAACTACGAGCTTTTATATTCCCTGCAGCATGTCTGTTTG**CAAGTTCTGCTCACGCGCTTCTGG**TCTCTGGAATACGCGTGTGGGACGGCTCAGGACAACGGTTC**CCCGCGGCGGTCCCTGCTAG**CCGGACACCGCTGGCAATCCCCCTGAAGTCCATAAACTCG**CCTTGTTCAGAACTTAAAGAGAAA**CAATCCTTGGTTGACATCGTTTATTGTCGTGGCATCCTAACTCAT**TCATTGAATGAATTGGCCATTT**TGTTATAATGAGCCGTCCGCCCTACGGAAACCTTGTTACGACTTCAT**CTTCCTCTAAATGATAAGATT**ATCCAAAGTGTTCACCGGACCAGAATGACAGCAAACTCGTTGTTGC**ATCGTTTACCGACTTTAG**ACGTTTCGGCAGTGGCTAGTATGCCACATAGTAAATTGCACACAAATG**TTGGCGGACAATTGCATGCGAT**ATGTTGGGCTTGTTAGAAAGGCCTGTTTGTCGAACGGTACTTGCA**CTGTACTACGCATTGTATAC**GCAACATGTGCACGCTGTTATTTGAAACGACTTGCTCATCAACTTT**CGATGGTACGCTACGTGCTTACCATGG**GACAACGGTTAACGGAGAATCAGTCCAGAAAAGTCTTCACTCTCCAGC**TCAGCTCCAGTTGAACAACAGCCTCCAA**AGCAACAATAATATCTAAATTCTAACGCAGGACTACCAGGGTATCTAATCCTGC**CGTATTACCGCGGCTGCT**TGCTGCCTCCCGTAGGACAACGCGAAGAACCTTACCAAAG**CTCTGACCGGTCTTTAATCGGA**ATTTAAGGTGGGCACTCTGGGCTTCCTCTCAGAACCCCTATCCA**CTAATGGAACGCATCCC**CGGACATGTGAACTCATGATCAGAAGTACAAACTCCTAAAAAACGTAGAGC**AATAACGATTTACGTGATGTAAC**GCTAATGGCTTGTTTATTGGTCATC

**Expected hits for all primers and probes on plasmid 3 (5’ to 3’)**

Cyclospora F: 60-38
Cyclospora P: 20-37
Cyclospora R: 1-18

Cystoisospora F: 61-84
Cystoisospora P: 86-109
Cystoisospora R: 110-129

Giardia F: 130-149
Giardia P: 151-170
Giardia R: 187-172

Entamoeba F: 256-235
Entamoeba P: 211-234
Entamoeba R: 188-209

E. histolytica F: 257-278
E. histolytica P: 280-301
E. histolytica R: 321-302

Blastocystis F: 391-370
Blastocystis P: 349-369
Blastocystis R: 322-347

Ancylostoma F: 392-414
Ancylostoma P: 416- 433
Ancylostoma R: 455-434

Ascaris F: 456-480
Ascaris P: 482-503
Ascaris R: 525-504

Necator F: 526-547
Necator P: 549-568
Necator R: 589-569

Trichuris F: 590-613
Trichuris P: 615-641
Trichuris R: 664-642

Strongy F: 665-688
Strongy P: 690-717
Strongy R: 746-718

16S F: 806-790
16S P: 772-789
16S R: 747-770

Lachno F: 807-828
Lachno P: 830-851
Lachno R: 872-852

Bacteroides F: 932-913
Bacteroides P: 896-912
Bacteroides R: 873-894

Crassphage F: 933-962
Crassphage P: 964-986
Crassphage R: 1011-987
